# Effects of line orientation in visual evoked potentials. Spatial dynamics, and gender differences of neural oblique effect

**DOI:** 10.1101/323782

**Authors:** Elena S. Mikhailova, Natalia Yu. Gerasimenko, Anna V. Slavutskaya

## Abstract

The aim of the study was to investigate neural mechanisms of orientation discrimination in humans by recording the 128-channel event-related potentials (ERPs) of the brain. Stimuli were black-and-white gratings having four orientations: horizontal, vertical, and oblique 45 and 135 deg. from vertical. Thirty-five healthy subjects (16 males, 19 females) participating in the experiments were asked to differentiate the orientations and push the button. The results showed that the orientation processing consists of two main stages. In the occipital, parietal and temporal areas the P100 and N150 components reflecting the early perceptual stage of processing showed amplitudes greater for oblique orientations than for cardinal ones - “inverse” oblique effect. Further, in the central-parietal and frontal areas the P300 and N400 components reflecting the later cognitive stage of processing showed the “classic” oblique effect with greater amplitudes for cardinal orientations. Females demonstrated lower ERP amplitudes and less pronounced differences between responses elicited by cardinal and oblique orientations. The results suggest that in humans the orientation discrimination occurs in a distributed neural network involving the early sensory as well as higher anterior cortical areas. The classic behavioral oblique effect might be controlled by later processing in the anterior cortical areas. The insufficient neural orientation selectivity observed in females may be considered as the biological basis of their poorer visuospatial abilities.

## INTRODUCTION

Visual perception critically depends on orientation-specific signals that arise early in visual processing. A large body of animal experiments has examined the mechanisms of orientation detection (Hubel, Wiesel, 1962; Li, Peterson, Freeman, 2003; Wang, Spelke, 2002). Despite the importance of orientation perception in human visual experience, the information about its neurophysiologic sources is limited. Of great significance was the discovery of the “oblique effect” by Campbell and Kulikowski (Campbell, Kulikowski, 1966). The oblique effect phenomenon denotes that the nervous system is more sensitive to stimuli of cardinal (vertical and horizontal) orientations than to oblique ones. The preference for cardinal orientations was later shown in the wide range of visual tasks (Appelle, 1972).

Human neuroimaging studies demonstrated only a moderate bias for cardinal orientations versus oblique ones (Aspell et al., 2010; Engel, 2000; Furmanski, Fink, 2001; Gur, 2000; Swisher et al., 2010). In the most cited study (Furmanski, Engel, 2000) the authors revealed a possible neurofunctional basis of this effect in humans by demonstrating that the activation of neurons in V1 is greater to horizontal and vertical orientations than to oblique (45 or 135) ones. This metabolic response was accompanied by the better detection of horizontal and vertical lines in comparison to oblique ones and lower thresholds of contrast sensitivity for cardinally oriented stimuli. However, there were other studies demonstrating diverse results. For example, Swisher and coauthors showed a higher neurometabolic response to oblique orientations (Swisher et al., 2010). Sun and coauthors discovered more voxels that preferred horizontal and vertical orientations in comparison with oblique ones, but they did not find any significant differences between cardinal and oblique orientations relying on the BOLD signal altitude (Sun et al., 2013). Some authors showed the priority of oblique orientations when testing oblique effect with natural scene content (Hansen, Essock, 2004), with broad-band noise stimuli (Yang et al., 2012) and during saccadic eye movements (Lee, Lee, 2008).

There is limited data concerning the response anisotropy for line orientation (unequal responses to cardinal and oblique orientations) in human event-related potentials (ERPs) (Arakawa et al., 2000; Moskowitz, Sokol, 1895; Proverbio et al., 2002; Takacs et al., 2013). Takacs and coauthors (Takacs et al., 2013) showed that oblique effect may also be present at the pre-attentive levels, and argued that the oblique effect is a fundamental phenomenon in visual perception. It should be emphasized that the ERPs’ data were mainly related to the relatively late (after 200 ms) stages of the information processing. The only study (Koelewijn et al., 2011) was aimed at analysing the early stage of the line orientation perception in the human visual cortex. Using MEG, these authors studied induced gamma and transient evoked responses to stationary circular grating patches of three orientations (0, 45, and 90 deg. from vertical). Surprisingly, they found that the sustained gamma response was larger for oblique than for cardinally oriented stimuli. This inverse oblique effect was also observed in the earliest (80 ms) evoked response, whereas later responses (latency 120 ms) showed a trend towards the reverse, “classic”, oblique response. This result was different from previous ERP (Moskowitz, Sokol, 1985) and fMRI (Furmanski, Engel, 2000) studies that showed only a single neural response that was greater for cardinal stimuli, compared to obliques. The authors explained this inconsistency in terms of the greater spatial resolution of MEG, compared to EEG, and its greater temporal resolution, compared to fMRI.

The important problem of gender specificity of orientation sensitivity has received little emphasis up to now, although numerous data point to the possibility of gender differences in the detection and coding of basic characteristics of visual space (Andersen et al., 2012; Collaer, Nelson, 2002; Collins, Kimura, 1997; Iachini et al., 2005; Goyette et al., 2012; Luyat et al., 2012).

Data relating to gender differences in the performance of visuospatial tasks are well represented in literature. Data relating to gender differences in the performance of visuospatial tasks are well represented in the literature. The gender differences were recorded during the three-dimensional mental rotation task (Collins, Kimura, 1997; Jordan et al., 2002; Tzuriel, Egozi, 2010), visual object construction test (Georgopoulos et al., 2001; Mikhailova et al., 2012), and navigation (Andersen et al., 2012). The differences in the performance of line orientation recognition task are described in a few behavioural studies (Caparelli-Dáquer et al., 2009; Collaer, Nelson, 2002).

In the recent study (Slavutskaia et al., 2014) we found that female performance was worse on the tasks requiring an accurate estimation of orientations in the Benton test (the determination of orientation of a short line by selecting it from a set of standard/etalon orientations) and an accurate identifying of the orientations of horizontal, vertical and oblique (45, and 135 degrees) lines. At the same time females and males did not differ when they had to determine the proximity of oblique lines to the horizontal, vertical and 45 degree lines. These data were assumed to be an indication of female insufficiency in metric abilities. The female insufficiency in some components of spatial cognition, in particular, in the encoding of the metric structure of the spatial relationships, was shown by Iachini and coauthors (Iachini et al., 2002). They found that while performing mental reproduction of the previously memorized spatial layout of objects males were more accurate in reproducing metric characteristics, namely, absolute and angular distances between objects. But males and females did not differ in recalling the position of objects. Furthermore, some authors demonstrate a female advantage for object location memory (Duff, Hampson 2001; Lejbak et al., 2009; Voyer et al., 2007). There is an opinion that females preferably adopt a non-metric strategy applied to individual objects (landmarks), whereas males’ metric strategy is applied to spatial relations (route) and spatial layouts as a whole (survey) (Coluccia, Louse, 2004).

In our previous study (Mikhailova et al., 2012) males and females revealed no differences in reaction time and accuracy at figure identification after its spatial transformation (displacement and rotation of the figure’s details). At the same time, there were significant differences in the image processing, and only males revealed the early perceptual stage sensitivity to the image transformation. The data probably imply different strategies of the visual processing used by males and females in visual-spatial tasks. One can assume that the gender differences in visual-spatial tasks performance may be related to differences in the ability to accurately identify the main spatial axes and deviations from them.

The goal of our present study was twofold: to fiurther investigate the neural mechanisms of orientation selectivity and to study how gender differences in the discrimination of line orientation are reflected in the ERPs.

## METHODS

### Subjects

The study involved forty-one subjects (21 females and 20 males) with normal vision. The average age was 23.3±0.9− and 22.2±0.8 for males and females, respectively. All experiments were performed with the consent for the study from the subjects (students of Biological faculty and faculty of Physics of Moscow State University, graduate students and young scientists of the Institute of High Nervous Activity and Neurophysiology of RAS) according to the protocol approved by the local ethics committee. Investigations were carried out in the first half of the day from 9 till 14 hours. The participation in the experiment was paid.

### Stimuli and procedure

ERP stimuli consisted of a stationary highcontrast black/white square-wave gratings of patches of four orientations (0°, 45°, 90° and 135° from horizontal, 5 cycle/degree, diameter 5.5 angular degree). The stimuli were presented in the centre of 17 inch Dell E1911c monitor (1280Ч1024 pixel resolution, 60 Hz refresh rate). An additional oval shield covered the vertical and horizontal edges of the monitor.

Stimulus duration was 100 ms, the interval between the stimuli randomly changed from 3 to 4 seconds. During this interval a grey fixation dot was continuously presented in the screen centre. Each stimulus was presented 34 times, and all session took approximately 25 min in total. There was a 5 minute break in the middle of the session. All displays were generated using E-Prime 2.0 software (Psychology Software Tools, Inc.). Before the session the participants were instructed which button corresponds to which particular grating’s orientation. The subjects were asked to identify the grating’s orientation and press the corresponding button on a Serial Response Box (Psychology Software Tools, Inc.). Subjects were instructed to maintain fixation for the entire experiment and to press a button as fast as possible at the termination of each stimulation period, to maintain attention. The individual accuracy (probability of correct answers in %) and RT (ms) were averaged separately for each orientation across all session. The trials involving incorrect or missing responses were excluded from the analysis of RT.

### EEG recording and signal processing

The EEG was acquired using a 128-channel system (*HydroCel Geodesic Sensor Net, Electrical Geodesics Inc., USA*), and all data were processed offline using the Net Station EEG Software. The impedance of all electrodes was kept below 50 kΏ during data acquisition. All electrodes were physically referenced to Cz (fixed by the EGI system) and then were re-referenced off-line to the averaged mastoid reference. The EEG was amplified with a band pass of 0.01–200 Hz, which was digitized on-line at a sampling rate of 500 Hz. During the off-line analysis of the EEG data, a finite impulse response bandpass filter with a low-pass frequency of 40 Hz was employed. The data were segmented relative to the test-stimulus onset (300 ms before and 700 ms after) and were sorted according to the stimuli orientation. Epochs contaminated by eye blinks, eye movements and muscle artifacts, as well as incorrect behavioral responses, were eliminated. The baseline began at 300 ms pre-stimulus and lasted for 300 ms.

ERP analysis23–25 clean trials were averaged for each stimuli orientation per participant. Mean amplitudes of four time windows surrounding the peak latency of distinct ERP components were included into our analysis: P100 (50–130 ms latency window), N150 (100–190 ms), and P300 (300–450 ms) peaks. In addition, based on the visual inspection, amplitudes in anterior N400 component (400–600 latency window) was analyzed. The amplitudes of the individual ERP components were measured as the average value of preselected eight right and in eight left groups of electrodes, covering occipital, posterior parietal, temporal scalp regions in posterior cortical areas, and more anterior central/parietal(anterior part) and frontal/prefrontal areas over the right and left hemispheres, correspondingly (Fig.1). The individual electrodes that were included in these groups were: 76, 83 (O_2_), 84 (right occipital group); 66, 70 (O_1_), 71 (left occipital); 90, 91, 96 (T_6_) (right temporal); 58(T_5_), 59, 65 (left temporal); 77, 85, 92(P_4_)(right parietal); 52(P_3_), 60, 67 (left parietal); 86, 93(near C_4_), 98 (right lateral parietal); 42 (near C_3_), 47, 53 (left lateral central); 78, 79, 87 (near C_4_) (right medial central); 37 (near C_3_), 54, 61 (left medial central); 2, 8,14 (right lateral frontal); 21, 25, 26 (left lateral frontal); 3, 9(FP_2_),10 (right medial frontal); 18, 22(FP_1_), 23 (left medial frontal); 4,122(F_8_), 123, 124 (F_4_)(right caudal frontal) and 19, 24(F_3_), 27(F_7_), 33(left caudal frontal).

**Figure 1.**
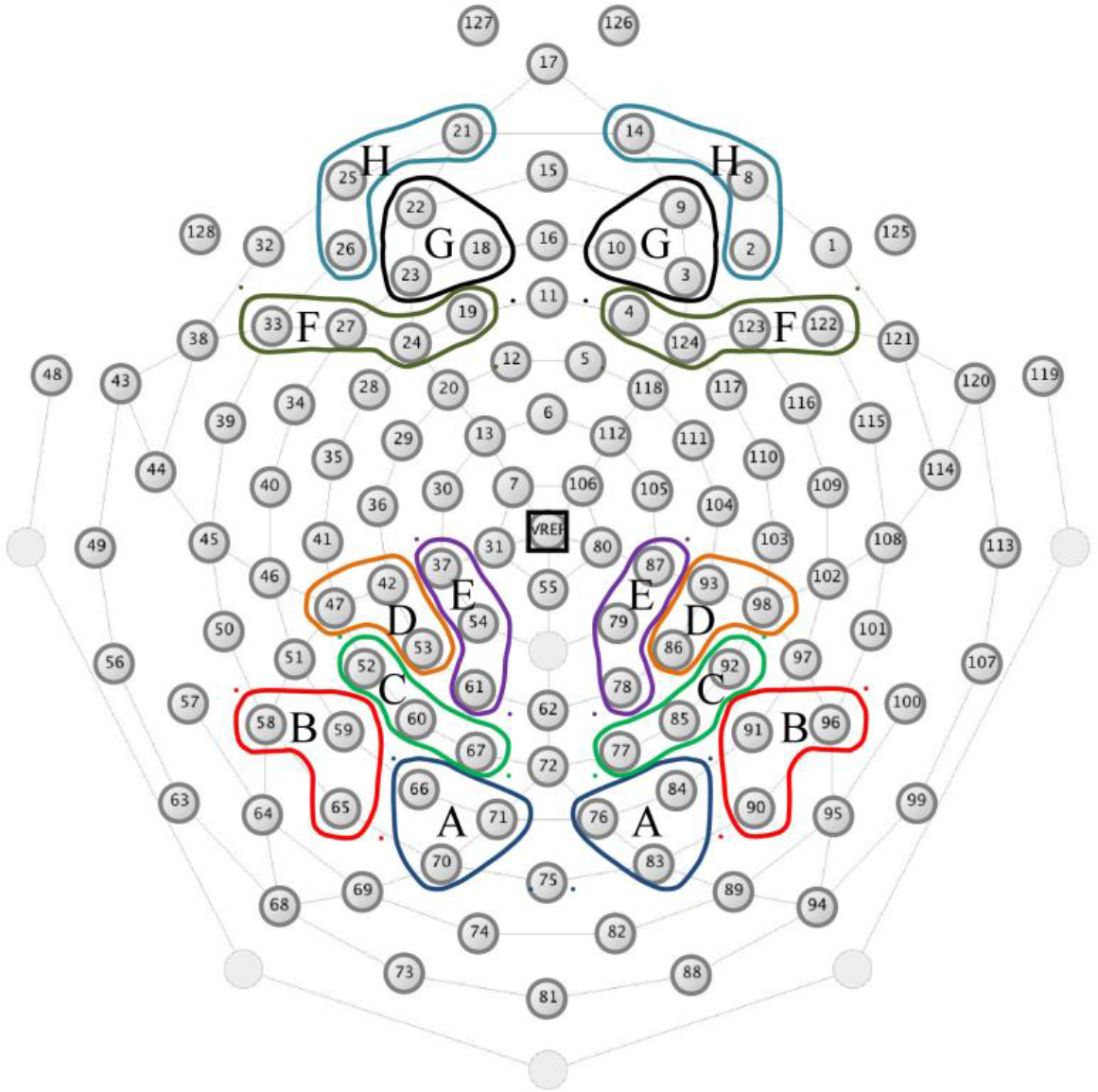
Arrangement of the high-density electrode arrays and the locations of the groups of electrodes over the skull. Electrode grouping for the symmetrical occipital groups (A) consisting of channels 66, 70 (O1), 71 on the left and 76, 83 (O2), 84 on the right, for the parietal groups (B) - channels 58(P7), 59, 65 on the left and 90, 91, 96 (P8) on the right, for the temporal groups (C) - channels 52 (P3), 60, 67 on the left and 77, 85, 92 (P4) on the right, for the central lateral groups (D) - channels 42, 47, 53 on the left and 86, 93, 98 on the right, for the central medial groups (E) - channels 37, 54, 61 on the left and 78, 79, 87 on the right, for the frontal caudal group (F) - channels 19, 24, 27, 33 on the left and 4, 122, 123, 124 on the right, for the frontal medial groups (G) - channels 18, 22 (FP1), 23 on the left and 3, 9 (FP2), 10 on the right, for the frontal lateral groups (H) - channels 21, 25, 26 on the left and 2, 8, 14 on the right.

The caudal components P100 and N150 were measured at the right and left occipital, temporal and parietal electrode groups, the component P300 was measured at the lateral and medial central electrode groups, and the anterior N400 component was measured at lateral, medial and caudal frontal groups. In the individual ERPs the amplitude extremums were measured within the corresponding time segments. The average amplitude value (adaptive maximum/minimum) for components P100 and N150 was counted within the 4-ms interval (2 ms before and 2 ms after the component peak), for components P300 and N400 - within the 8-ms interval (4 ms before and 4 ms after the component peak).

The statistical analysis of the amplitude values of the ERP components was executed with the help of ANOVA RM variance analysis (resetting method). The ORIENTATION, HEMISPHERE, REGION (group of electrodes) factors were used as within-subject factors and GENDER - as the between-subject factor. The Greenhouse-Geisser correction for non sphericity was applied whenever appropriate. The Newman-Keuls correction for the multiple comparisons was taken into account at the post-hoc horizontal (0 deg.), vertical (90 deg.), and oblique 45 deg and 135 deg. orientations.

## RESULTS

### Behavioral data

A higher accuracy and a lesser RT were shown for the identification of horizontal and vertical orientations in comparison with oblique ones. The ANOVA RM variance analysis of the accuracy and the RT was conducted. It included Orientation (horizontal, vertical, oblique 45 deg., oblique 135 deg.) and Gender (males, females) factors. Significant effects of Orientation (F3,117=6.62; p<0.001) were revealed for the accuracy. Post-hoc comparisons showed that the accuracy of identification of cardinal and oblique orientations significantly differs only in the female group: 45 deg. and 135 deg. orientations were determined worse than cardinal ones (0.002 < p <0.02). The significant effects of Orientation (F3,117=6.62; p<0.001) and Gender (F1,38=4.28; p<0.05) were revealed for the RT. Post-hoc comparisons showed that both male and female groups have the RT, higher for oblique orientations (p <0.0001) in comparison with cardinal ones.

### ERP data

After a visual inspection five subjects (four males and one female) were excluded from the analysis due to a large number of artifacts in their EEG recordings. Thus, the final ERP analysis included the data on 35 subjects (16 males and 19 females).

Fig. 2 and Fig.3 display grand average ERP waveforms elicited by cardinally and obliquely oriented gratings in males and females, respectively. The observed components were named according to their latencies, polarities and topographic properties as the P100 and N150 with maximum over the caudal areas, P300 with maximum over the parietal and central areas, the N400 with maximum over the frontal areas. We measured and analyzed the P100 and N150 components at occipital, temporal and posterior parietal electrode groups, P300 component at anterior parietal and central electrode groups and N400 component at frontal and prefrontal electrode groups.

**Figure 2.**
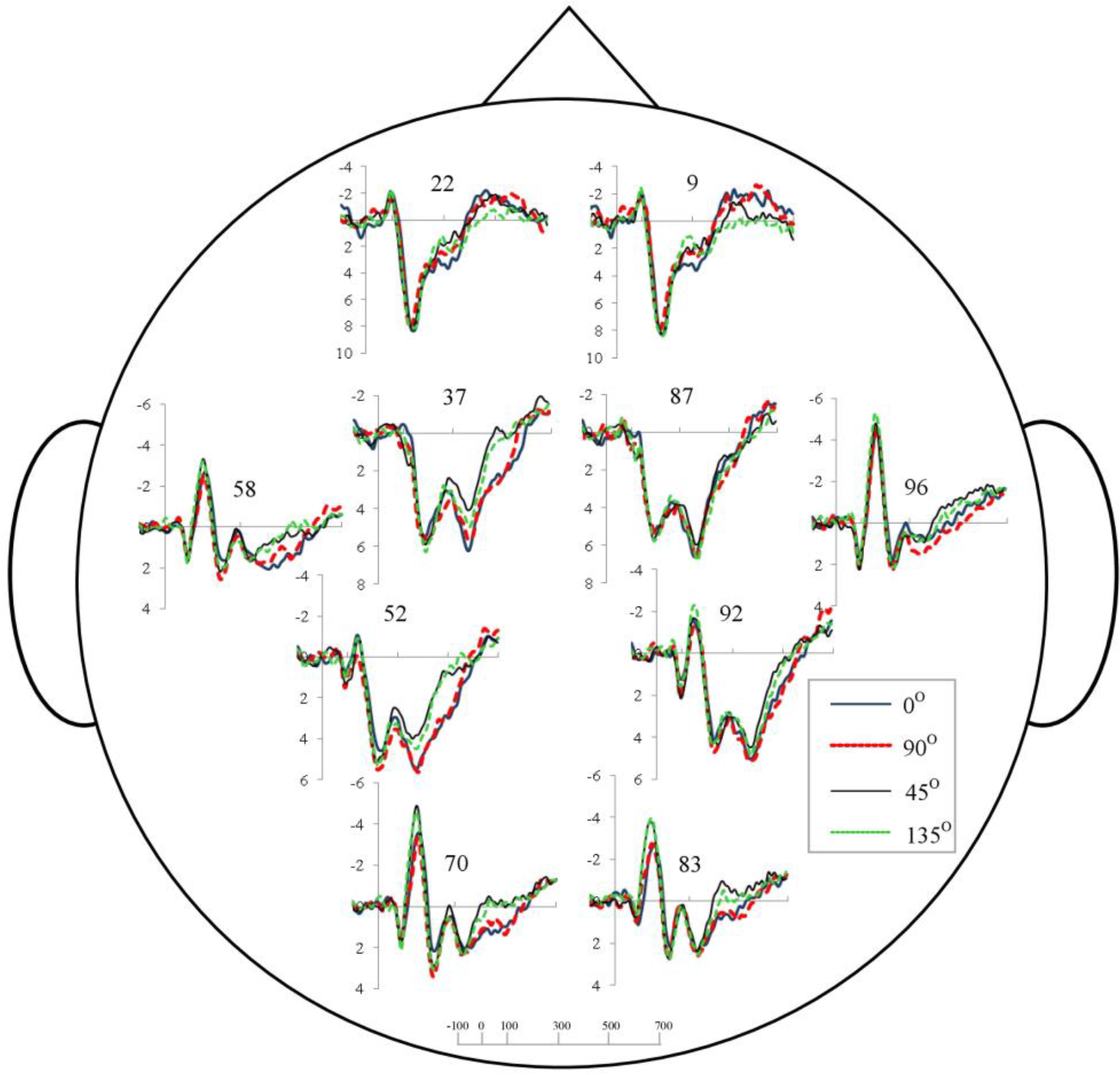
Grand average ERP waveforms for the horizontal (0°), vertical (90°) and oblique (45° and 135°) gratings at the occipital (channels 70 and 83), parietal (channels 52 and 92), temporal (channels 58 and 96), central (channels 37 and 87) and frontal (channels 9 and 22) electrodes in male group. Vertical scale represents voltage amplitude in μV and horizontal scale displays latency in ms.

**Figure 3.**
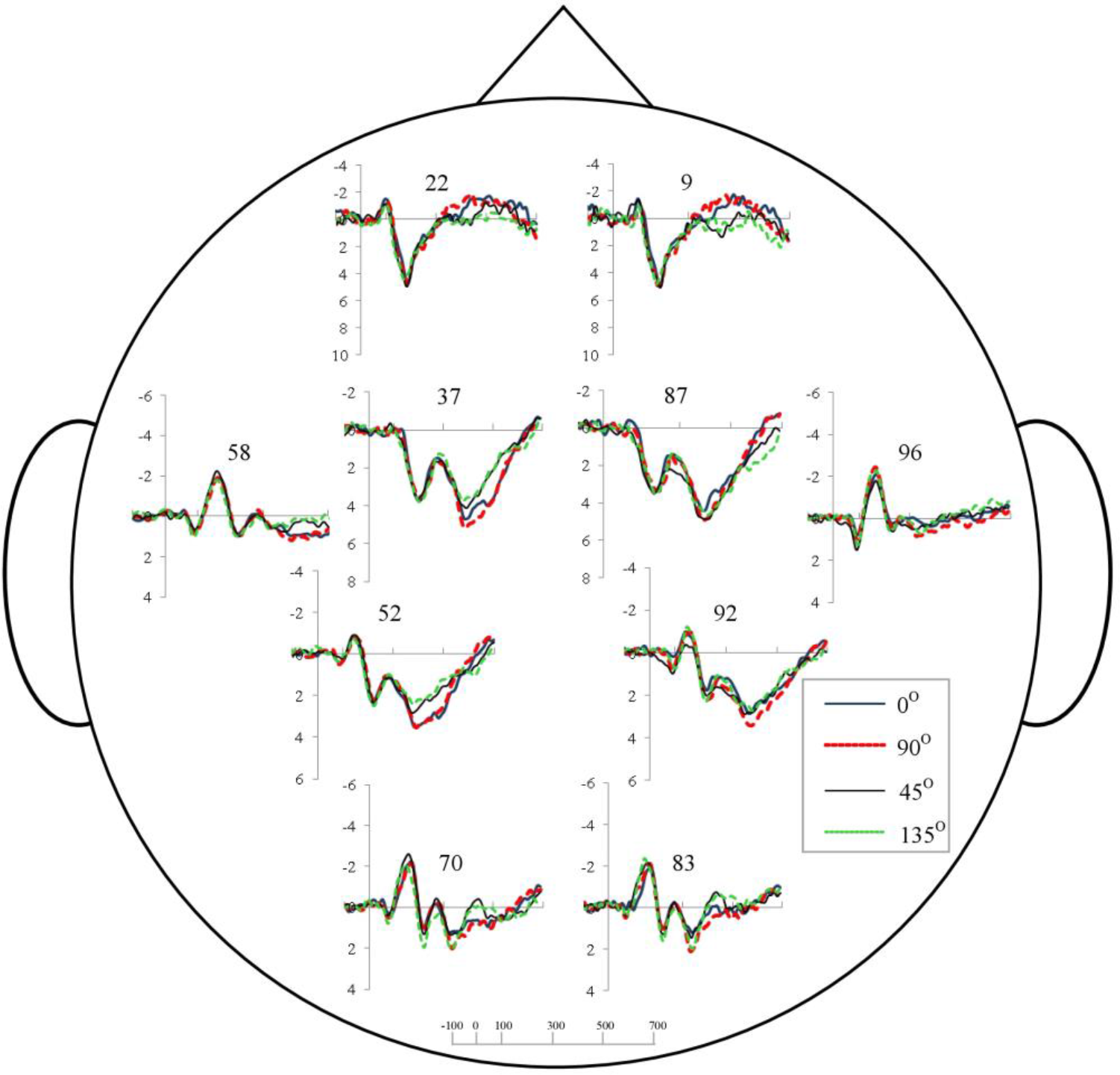
Grand average ERP waveforms for horizontal (0°), vertical (90°) and oblique (45° and 135°) gratings at the occipital (channels 70 and 83), parietal (channels 52 and 92), temporal (channels 58 and 96), central (channels 37 and 87) and frontal (channels 9 and 22) electrodes in female group. Vertical scale represents voltage amplitude in μV and horizontal scale displays latency in ms.

### P100 component

The amplitude of P100 was analyzed in three electrode groups: occipital, temporal and parietal ones which later will be labeled as Regions. Overall ANOVA RM [Orientation × Region × Hemisphere × Gender] showed a significant interaction Orientation × Region (F6,186=2.63; p<0.03). This result indicates that the responses evoked by different orientations depended on the cortical site. To find the region specificity of the effect of Orientation, the further analysis was carried out for the occipital, temporal and parietal regions separately.

In the parietal region a significant main effect of Orientation (F3,93=2.88; p<0.05) and a significant interaction Orientation × Hemisphere × Gender (F3,93=2.98; p<0.04) were found. The grand averaged ERPs in Fig.2 and Fig.3 and the diagrams in Fig.4 illustrated, that oblique 45 deg. orientation elicited the larger P100 amplitude than the latter elicited by cardinal (horizontal) one. What is the most important, the post-hoc comparisons found significant differences in the left parietal area at the contrasts horizontal *vs* 45 deg. (p<0.003) and at horizontal *vs* 135 deg. (p<0.03) only in males.

**Figure 4.**
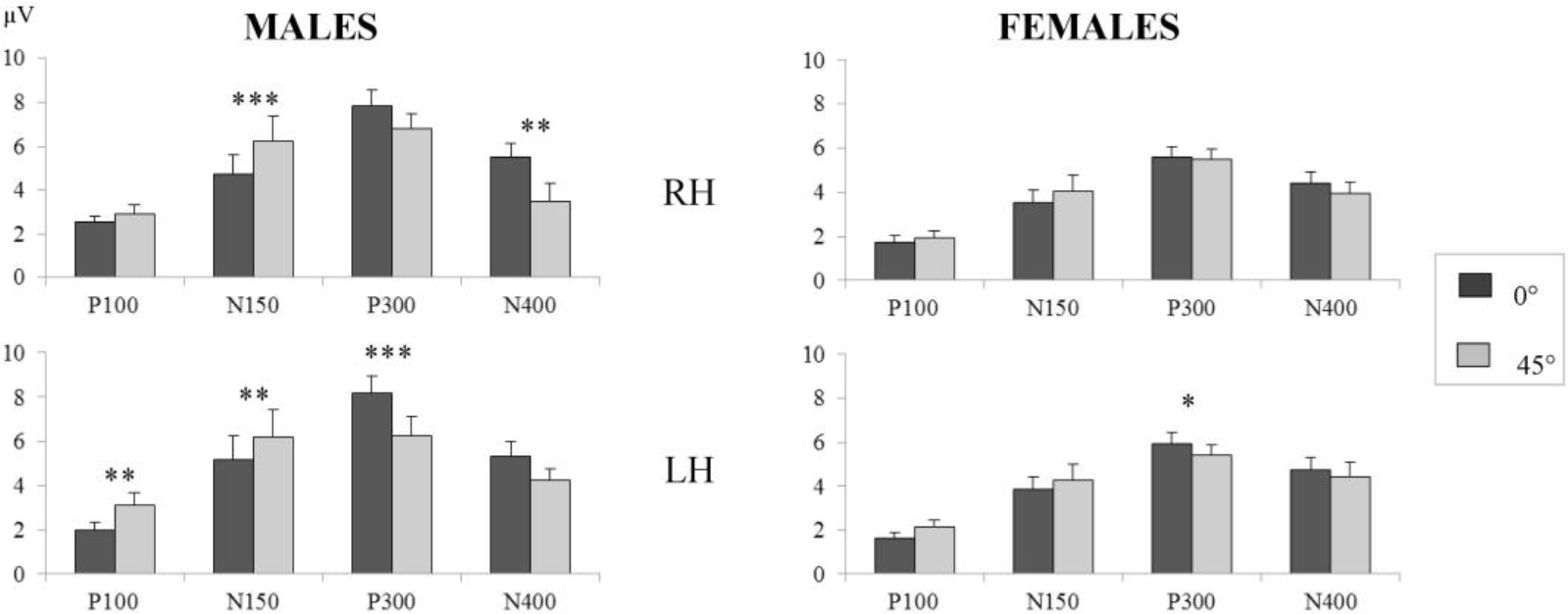
Mean amplitudes of ERP components for horizontal (0°) and oblique (45°) gratings in the right and left hemispheres in male and female groups. The bars represent average mean amplitude of components over groups of electrodes at parietal (for P100), occipital (for N150), central medial (for P300) and frontal lateral (for N400) recording regions in the left and right hemispheres. Asterisks *, ** and *** represent significance levels p < 0.05 p < 0.01 and p < 0.001, respectively. Errors bars indicate SEM.

**Figure 5.**
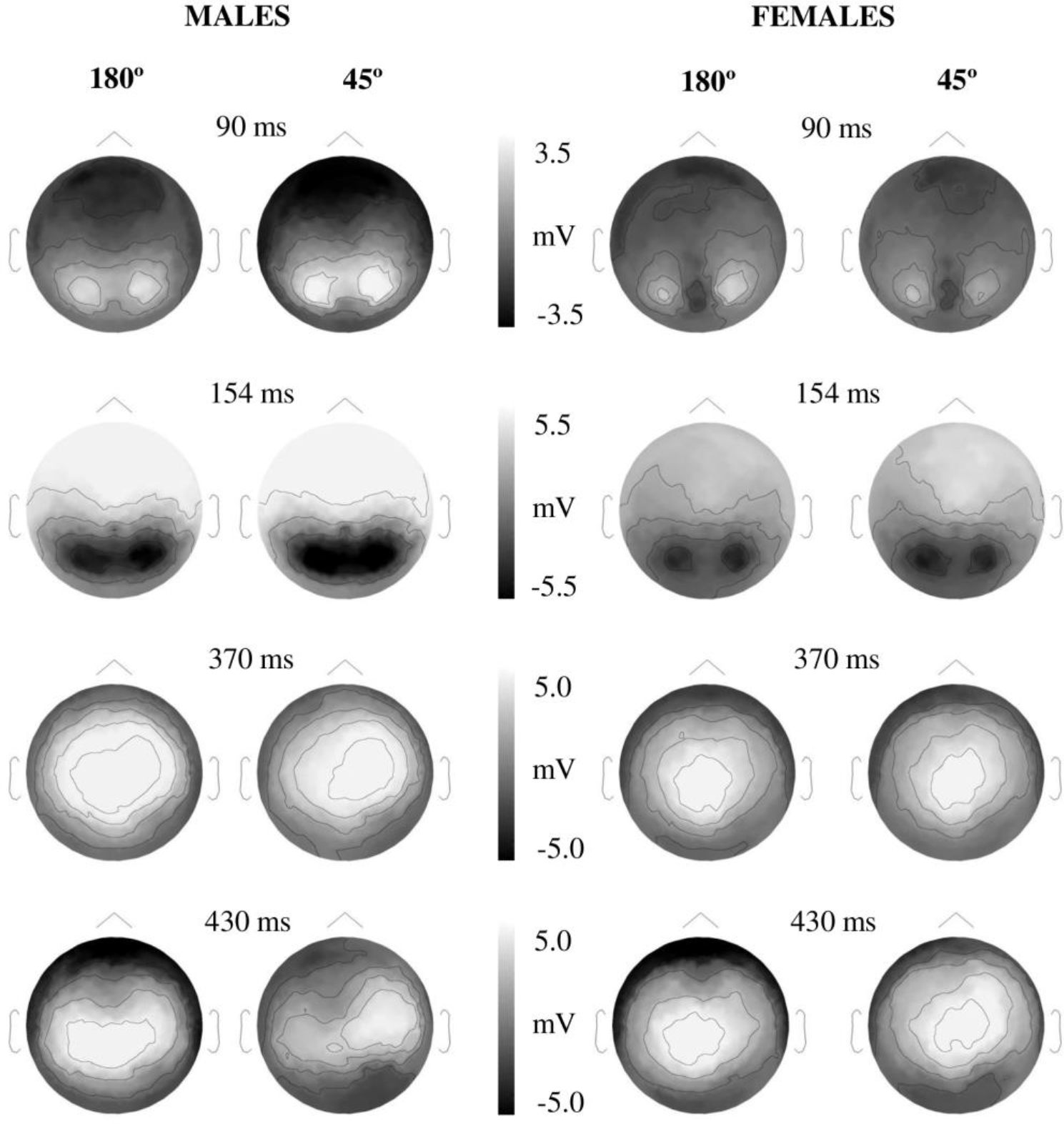
Topographical representations of brain activity evoked by horizontal (0°) and oblique (45°) gratings at the moments of the P100 (90 ms), N150 (154 ms), P300 (370 ms) and N400 (430 ms) peaks in groups of males and females.

In the occipital region there were no significant main effects of Orientation, Hemisphere and Gender. There was found an interaction Orientation × Hemisphere × Gender (F3,93=3.08; p<0.05), but the further post-hoc comparisons didn’t show any significant differences.

In the temporal region no significant main effects and interactions were observed.

### N150 component

The amplitude of N150 was analyzed in three regions: occipital, temporal, parietal ones. Overall ANOVA RM [Orientation × Region × Hemisphere × Gender] showed significant main effects of Orientation (F3,99=5.41; p<0.005) and Region (F2,66=20.291; p<0.0005). Also we found significant interactions Orientation × Region (F6,198=5.45; p<0.001) and Orientation × Hemisphere (F3,99=4.74; p<0.01). Further analysis was carried out for every region.

In the occipital region we found more expressed significant main effect of Orientation (F3,99=8.36; p<0.0005). Also, there was a significant interaction Orientation × Gender (F3,99=3.63; p<0.02). The grand averaged ERPs in Fig. 2 and Fig. 3 and the diagrams in Fig. 4 illustrated, that oblique 45 deg. orientation elicited the larger N150 amplitude than the horizontal one. Further post-hoc comparisons confirmed the significance of these differences in the right (0.002<p<0.0002) and in the left (0.05<p<0.001) hemispheres only in males

In the temporal region, as in the occipital one, a main effect of Orientation was found (F3,99=5.70; p<0.003). Also a significant main effect of Gender (F1,33=3.90; p<0.05) was found, and the amplitude N150 is higher in males compared with females. Similar to the occipital region, males and females revealed different effects of Orientation, that were expressed in a significant interaction Orientation × Gender (F3,99=5.05; p<0.005). More specifically males exhibited the marked differences in the responses to cardinal and oblique orientations. Further post - hoc evaluation found significant differences at the contrasts horizontal *vs* oblique (0.05<p<0.005), vertical *vs* oblique (0.05<p<0.005) in the right hemisphere, and vertical *vs* oblique (p<0.005) in the left one.

In the parietal region no significant effects of Orientation and Gender were observed.

### P300 component

The amplitude of P300 was analyzed in the medial and the lateral central electrode groups. Overall ANOVA RM [Orientation × Region × Hemisphere × Gender] showed significant main effects of Orientation (F3,96=6.81; p<0.001), Region (F1,32=129.3; p<0.00005), and Gender (F1,32=6.33; p<0.02). Also, there was a significant interaction Orientation × Hemisphere (F3,96=9.18; p<0.00005). Further analysis was carried out for every region.

In the medial central groups, a significant main effect of Orientation (F3,96=7.28; p<0.0001), and an interaction Orientation × Hemisphere (F3,96=8.24; p<0.0001) were found. What is the most important, P300 amplitude revealed the orientation bias which was different from the for P100 and N150 orientation bias. Figure 4 shows, that the P300 amplitude in responses to cardinal orientations was higher than to oblique ones. Significant differences were found in the left hemisphere only. In males a post - hoc comparison found the differences at the contrasts horizontal *vs* oblique 45 deg. (p<0.0005), vertical vs oblique 45 deg. (p<0.0001) and vertical *vs* oblique 135 deg. (p<0.005). In females the orientation differences were weaker and were found at the contrasts horizontal vs oblique 45 deg. (p<0.02) and vertical vs oblique 45 deg. (p<0.02). A significant main effect of Gender was found (F1,32=7.39; p<0.01), and P300 amplitude was higher in males than in females (Fig.2–4).

In the lateral central group as in the medial one, a significant main effect of Orientation was found (F3,96=7.27; p<0.0001). Cardinal orientations elicited the larger P300 amplitude than oblique ones (Fig. 4). In addition, a significant main effect of Gender (F1,32=6.72; p<0.01) was found, and the P300 amplitude was higher in males comparing with females (Fig. 2–4). A significant interaction Orientation × Hemisphere was found (F3,96=6.52; p<0.001). Separate post-hoc comparisons showed, that in males the P300 amplitude was higher in responses to cardinal orientations than to oblique ones (0.0001<p<0.01), but only in the left hemisphere (Fig. 4). In females the differences between cardinal and oblique orientations were weaker (p<0.02) and were found in the left hemisphere only.

### N400 component

The amplitude of N400 was analyzed in the three frontal electrode groups: lateral, medial and caudal ones (Fig. 1). Overall ANOVA RM [Orientation × Hemisphere × Region (lateral, medial, caudal) × Gender] showed significant main effects of Orientation (F3,96=4.66; p<0.01), and Region (F2,64=29.97; p<0.0001). Moreover, there were significant interactions Orientation x Region (F6,192=3.38; p<0.01), indicating that orientation-dependent reactions are not the same at the three different frontal regions. The interactions Orientation × Hemisphere × Gender (F6,96=2.95; p<0.05) and Orientation × Region × Gender (F6,96=2.52; p<0.05) supposed, that the effect of orientation is different in males and females. Further analysis was carried out for every region.

In the more anterior frontal lateral region a significant main effect of Orientation (F3,96=7.74; p<0.001) was found. As illustrated by averaged ERPs (Fig. 2–3) and graphs (Fig.4) amplitude N400 is higher in responses to cardinally oriented gratings.

Further post - hoc evaluation found the significant differences between cardinally and obliquely oriented gratings only in males at the contrasts horizontal *vs* oblique 45 deg. (p<0.006), horizontal *vs* oblique 135 deg. (p<0.0003), and vertical *vs* oblique 135 deg. (p<0.01) in the right hemisphere (Fig.4).

In the medial and caudal frontal regions the effect of Orientation was less pronounced. In the medial region a main effect of Orientation (F3,96=3.13; p<0.05) and an interaction Orientation × Hemisphere (F3,96=4.14; p<0.01) were found. In the caudal region an interaction Orientation x Hemisphere (F3,96=6.45; p<0.001), and a main effect of Hemisphere (F1,32=5.57; p<0.02) were found. Similar to the lateral region, in the medial and caudal frontal regions a post - hoc evaluation showed the differences at the contrasts cardinally oriented gratings *vs* obliquely oriented gratings (0.01<p<0.05) in the right hemisphere.

Further post - hoc evaluation found the significant differences between cardinally and obliquely oriented gratings only in males. They were found at the contrasts horizontal *vs* oblique 45 deg. (p<0.02) and horizontal *vs* oblique 135 deg. (p<0.01) in the right medial region, and at the contrasts vertical *vs* oblique 135 deg. (p<0.01) and horizontal *vs* oblique 135 deg. (p<0.05) in the right caudal region. Females did not show any significant differences between orientations. There was only tendency at the contrasts horizontal *vs* oblique 45 deg. and horizontal *vs* oblique 135 deg. (p=0.08) in the left anterior region, and at the contrasts horizontal *vs* oblique 45 deg., horizontal *vs* oblique 45 deg. (p=0.08) in the right medial and caudal regions.

## DISCUSSION

The goal of current study was to determine whether peak amplitudes of early (P100 and N150) and later (P300 and N400) ERP components differed when evoked by gratings containing cardinal and oblique orientations, and whether the found orientation bias differed at earlier and later stages of visual processing. Thus, we studied the dynamics and topography of cortical evoked responses to cardinal and oblique orientations throughout sensory and cognitive stages of visual processing. We also focused on the gender differences in visual processing of orientations. It was hypothesized that gender differences in orientation selectivity might/could underlie the general differences between males and females in visual-spatial tasks performance.

We revealed orientation-selective activation during the P100 and N150 components in the posterior cortex, which reflects the differences in the early sensory processing of cardinal and oblique orientations. In this time window we unexpectedly found the inverse oblique effect which appeared in the greater response to oblique over cardinal orientations. A similar inverted neural oblique effect was described by Koelewijn and coauthors (Koelewijn et al., 2011). Using MEG, they found the inverse oblique effect in the earliest (80 ms) evoked response over parietal and occipital sensors. The inverse oblique effect was also shown in several other studies (Hansen et al., 2010; Lee, Lee, 2008; Swisher et al., 2010; Yang et al., 2013;).

The origin of the inverted oblique effect remains unclear till now. Koelewijn and coauthors assumed that the stronger response to oblique stimuli might reflect increased tuning widths of visual cortex neurons for these orientations (Koelewijn et al., 2011). The results of some studies evidenced the more complex neural networks for oblique orientations processing. Edden and coauthors showed that behavioral orientation detection thresholds were negatively correlated with visual cortex GABA concentration and with gamma oscillation frequency for obliquely oriented patterns but not for vertically oriented one (Lejbak et al., 2009). These authors suggested that the difference between cardinal and oblique performance may not arise from primary visual cortex itself, but may result from top-down modulation from higher extrastriate visual areas. Another possible explanation of the high amplitude response to oblique orientations is based on the increased visual attention when participants perform a more difficult task of the oblique orientation recognition. It is possible the classic oblique effect could not have specific ERP manifestations in the visual areas, and early ERP expressions of the oblique effect depend to a large extent on the experimental conditions such as the task performed.

In our study the earlier orientation selective response (P100) was found in the parietal cortex. The parietal cortex is known to be critical for a lot of visual-spatial functions including eye movements, reaching movements, stimulus motion, and spatial attention (Andersen et al., 1997; Lauritzen et al., 2009; Luyat et al., 2005; Luyat, 2008; Silver, Kastner, 2009). The posterior parietal cortex is known to be essential for the representation of space and for the transformation of spatial information into a motor coordinate framework (Andersen, Buneo, 2002). Several studies demonstrated that the parietal cortex in animals and in humans plays a very important role in the organization of the multimodal reference frame (Fasold et al., 2002; Luyat et al., 2005). It is also well known that the processes in the posterior parietal cortex are critical for spatial working memory, namely for the maintenance of spatial information and topographic maps of visual space (Curtis, 2006).

Contrary to our expectations, we did not find the P100 orientation-selective answer in the occipital area. But in our recent study using the dipole modeling (wMNE) of the ERPs components we observed significant source current density differences between P100 evoked by cardinal and oblique orientations in *superior, middle and inferior occipital gyri, calcarine fissure*, and *lingual gyrus* (Krylova et al., 2015).

The orientation effect for the N150 was the greater than for the P100. It was observed in the occipital and temporal areas. Similar to the P100, the N150 amplitude demonstrated the inverted oblique effect and the N150 was higher for oblique orientations than for cardinal ones. The importance of N150 time window for the oblique orientation discrimination is emphasized in some studies. According to the investigation of Takacs and coauthors, in the occipital and parietal areas the negativity within 120–190 ms time window reflected an inclination from the cardinal axes, and the higher the negativity amplitude with the larger inclination (Takacs et al., 2013). The correlation between the negative deflection within this time window and the oblique orientation processing was confirmed by other authors (Song et al., 2010). In this study short-term training improved oblique orientation discrimination simultaneously with the N1b (latency of 170 ms) amplitude decreasing at lateral occipito-temporal channels. In was also found that the deviancy from cardinal orientation, which equals to the difference between the given orientation and closest cardinal orientation, demonstrated the strong correlation with the N1b amplitude in the posterior leads.

In our study the orientation selective responses in the N150 window were found in the temporal region. The N150 amplitude was greater to oblique orientations compared with answers to cardinal ones. Some authors also found orientation selectivity for temporal cortex responses. Using optical imaging, Xu and coauthors discovered that the middle temporal visual area (a visual area highly sensitive to moving stimuli) of the monkey *Aotus trivirgatus* was more devoted to representing cardinal than oblique orientations, and that the anisotropy was more prominent in parts representing central vision (Xu et al., 2006). In comparison, an overrepresentation of cardinal orientations in the representation of central vision in monkey V1 was relatively small and inconsistent. The authors supposed that the data could explain the greater sensitivity to motion discrimination when stimuli are moved along cardinal meridians. The orientation-selective answers in the temporal cortex were also described in humans in the P300 time window (Proverbio et al., 2000).

Supposedly the different N150 answers to oblique and cardinal orientations may reflect the basis of the early processing of different orientations. These results are consistent with the classical notion that the “N1” wave (140–200 ms) reflect the operation of a discrimination process (Luck, Hillyard, 1995) and focused in regions of inferior occipital-temporal cortex (Hopf et al., 2002).

The later components P300 and the N400 reflect the later stage of information processing in the identification of line orientation. The amplitudes of these components were assessed over the central-parietal (P300) and frontal (N400) electrode groups.

As a general remark, it should be noted that the P300 and N400 showed the classic oblique effect with the prevalence of cardinal orientations that was different from the inverted oblique effect at the first stage of orientation processing. The classic oblique effect revealed in the P300 and N400 correlated with the behavioral predominance of the basic orientations.

When discussing the P300, it should be pointed out that our results are in line with the conclusion that the central-parietal P300 component was modulated by the meaning and significance of the stimulus (for review see Ferrari et al., 2010).

Actually, vertical and horizontal orientations that correspond to the fundamental spatial axes evoke the more prominent P300. Consequently, under our experimental conditions the environmental significance of line orientation may be considered as a critical factor modulating the P300 amplitude. Our results are also consistent with a large number of studies that emphasize the relationship between the P300 and information content and also show that the P300 amplitude may reflect the processing of informationally rich events (Bledowski et al., 2004; Donchin et al., 1973; Kok, 2000).

In our present study the orientation differences found in the central-parietal area were statistically significant both in the medial and the lateral regions of this area. These findings are consistent with the ERP data according to which the classical P300 component or P3b occurs at 300–600 ms after a target stimulus and has a widespread parietal distribution on the scalp (Donchin et al., 1973; Donchin, Coles, 1988). When studying P300 generators in visual target and distractor processing, Bledowski with coauthors found the P300 sources both in the parietal regions (inferior parietal lobe - IPL and posterior parietal cortex - PPC) and in the inferior temporal (IT) cortex (Bledowski et al., 2004). They suggested that the IT source activity reflects the categorization process of the visual stimuli, whereas the generators in the parietal region reflect the stimulus-driven (exactly IPL) and top-down (exactly PPC) attentional processes that modulate and control the event categorization.

We showed that the P300 differences between orientations were statistically significant only in the left hemisphere. The data are consistent with the previous study of Proverbioa and coauthors reporting that an anterior P300 to target oriented gratings at central-parietal sites was of a greater amplitude in the left hemisphere than in the right one (Proverbio et al., 2000). Although these authors did not compare the P300 responses to vertical and oblique orientations, the reported data show more prominent orientation differences in the left hemisphere. We believe that our results are in line with the hypothesis that there is a special involvement of the left hemisphere in object discrimination (Georgopoulos et al., 2001; Proverbio et al., 2000).

The N400 as well as the P300 revealed the classic oblique effect, although the N400 differences between orientations were less expressive than the P300 ones. It is known that the frontal negativity N400 reflects later cognitive processing of a wide range of stimuli or tasks (for review see Kutas, Federmeier, 2011; Schendan, Lucia, 2010). Some authors consider the N400 as a modality dependent but not modality-specific electrophysiological marker of processing in a distributed semantic memory system (Kutas, Federmeier, 2011). Sitnikova and coauthors suggested that comprehension of the visual real world might be mediated by two neurophysiologically distinct semantic integration mechanisms (Sitnikova et al., 2008). One of these mechanisms, reflected by the anterior N400-like negativity, maps the incoming information onto the connections of various strengths between concepts in semantic memory. Hence, in our study the higher frontal N400 evoked by basic orientations might reflect their easier extraction from semantic memory.

When discussing the frontal cortex orientation-selective responses it is necessary to highlight the numerous data, showing the prefrontal cortex, specifically dorsolateral prefrontal cortex (dlPFC), contribution to processing spatial information and particularly to encoding cue locations (Burgess, 2006; Ma et al., 2003; Ma et al., 2012; Yang et al., 2012). In particular, dlPFC provides an egocentric mechanism that represents the location of an object with respect to the observers’ head (Burgess, 2006; Ma et al., 2003). In our current study we analyzed the N400 amplitude over three frontal electrode groups: lateral, medial and caudal. ANOVA RM showed that orientation-dependent answers were different in these electrode groups. The most significant orientation effect, that manifested itself in the higher N400 in response to cardinally orienting gratings, was found in the lateral frontal electrode group, apparently located over dlPFC. Taking into account these data, we suppose that the orientation selectivity of the N400 may be related to the frontal spatial frame system mechanism.

In our current study we observed clear gender differences in the orientation discrimination performance and also in the neural answers. The findings support our hypothesis that gender differences in visual-spatial tasks performance might be related to differences in the ability to recognize the basic orientations and deviations from them. The higher P100 and N150 amplitudes found in the occipital, temporal and parietal regions in males might indicate a higher activity during the sensory stage of orientation processing, providing more effective early orientation discrimination. This finding is consistent with the results of Gur and coauthors (Gur et al., 2000). While examining gender differences in BOLD answer for the judgment of line orientation task they recorded more extensive activation in posterior cortical areas in males than in females. Recently, using the modeling of the distributed intracranial dipole sources of the P100 and N150 by the weighted Minimum Norm Estimates, we found that males and females engage similar neural populations in the early orientation processing, but males show higher source current density in the occipital, parietal and temporal areas in the orientation discrimination task (Krylova et al., 2015).

During the sensory stage of the orientation discrimination there was another difference between males and females. The early ERP components demonstrated significant interactions Orientation × Gender. They were observed in the fact that the differences between orientations were more distinct in males than in females, especially in the N150 time window. In the first ‘initial classification’ state before 200ms, initial feedforward activation of sensory areas supports feature detection, structural encoding, and perceptual categorization (Bledowski et al., 2004; Hopf et al., 2002; Luck, Hillyard, 1995). This result shows the gender dependence of the early processing of line orientations. We believe that the females have an insufficient neural mechanism responsible for the early orientation discrimination. This may be an important biological factor determining the gender differences in orientation discrimination performance.

Similar to the sensory stage, during the later period 200–500 ms males demonstrated greater P300 and N400 responses than females. These differences localize in the central-parietal and frontal areas and reflect the gender specificity of the cognitive stage of orientation processing. This stage integrates various simultaneous brain processes such as allocation of attentional resources, estimation of stimulus probability and significance, stimuli congruency, activation of visual object knowledge for a category decision, and cognitive decisions about visual objects (Ferrari et al., 2010; Kutas, Federmeier, 2011; Schendan, Lucia, 2010).

The obtained results demonstrate that females process cardinal and oblique orientations differently than males. It is noteworthy that in the orientation discrimination task it is not only the sensory but also the cognitive stages are dependent on gender. Our study appears to be the first one to demonstrate a physiological correlate for gender differences in the processing of cardinal and oblique orientations.

## CONCLUSION

The findings presented here indicate that the orientation processing consists of two main stages - sensory and cognitive ones. The initial sensory stage of processing engages early visual areas. This stage is characterized by the inverted oblique effect manifested in neural responses which are greater to oblique orientations than to cardinal ones. The next cognitive stage engages central-parietal and frontal-prefrontal areas and reveals the classic oblique effect with greater neural responses to cardinal orientations. The orientation-selective responses found in more anterior areas may indicate the importance of orientation perception for planning goal directed spatial behavior. Females demonstrated lower ERP amplitudes and less pronounced differences between responses elicited by cardinal and oblique orientations. This insufficient neural orientation selectivity may be considered as the biological basis of females’ poorer visuospatial abilities.

## Acknowledgments

*This research was funded by Russian Academy of Sciences)*.

